# A data-driven rediscovery of the specificity-conferring code of adenylation domains in nonribosomal peptide synthetases

**DOI:** 10.64898/2026.06.15.732251

**Authors:** Zhengjian Li, Kenan A.J. Bozhüyük, Olga V Kalinina, Dietrich Klakow

## Abstract

Nonribosomal peptide synthetases (NRPSs) are large modular enzymes that assemble structurally diverse peptides, many of pharmacological importance, including antibiotics and immunosuppressants. Within each NRPS module, the adenylation (A) domain selects the substrate to be incorporated, a choice governed by a small set of residues lining the binding pocket. For two decades, computational prediction of A-domain substrate specificity has relied on residue sets—most prominently the Stachelhaus code and the 34-residue “8 Å code”—that were defined by spatial proximity to the substrate rather than by demonstrated predictive value. Here we revisit which residues govern substrate specificity from a purely data-driven perspective. We assembled a non-redundant dataset of 5,366 A-domain sequences (4,693 bacterial and 673 fungal) and used information-theoretic measures to rank alignment positions by their statistical association with substrate identity, without restricting candidate positions to any predefined structural shell. This procedure yielded two compact, kingdom-specific codes: IG15B (15 positions) for bacterial and IG13F (13 positions) for fungal A-domains. Both match or exceed the predictive accuracy of the 34-residue 8 Å code while using fewer than half its positions, and both independently recover the majority of the classical Stachelhaus positions. Notably, our analysis identifies four positions (242, 280, 281, and 284) that lie outside all conventional codes yet carry non-redundant specificity information and co-localize with classical determinants on two helices flanking the binding pocket. These positions provide new candidate sites for the rational engineering of A-domain specificity.

**Author summary:** Many clinically important drugs—including antibiotics such as vancomycin and immunosuppressants such as cyclosporin—are nonribosomal peptides, assembled by large enzymes known as nonribosomal peptide synthetases. These enzymes contain adenylation domains that act as molecular gatekeepers, each selecting one chemical building block to add to a growing peptide. Identifying which amino acids within a domain determine this choice is central both to predicting what an enzyme produces and to re-engineering it to make new compounds. For over twenty years, researchers have approached this question by selecting the amino acids that sit physically closest to the substrate. However, being close to the substrate does not guarantee that a residue actually influences substrate selection. In this work, we instead let the data decide: using thousands of adenylation domain sequences, we measured which positions are statistically most informative about the substrate, using information gain, mutual information and *χ*^2^ statistic. We found that far fewer positions than conventionally used are sufficient to predict specificity, and—importantly—we identified several influential positions that earlier approaches had overlooked because they lie just beyond the conventional distance cutoff. These positions offer promising new targets for engineering these enzymes to produce novel peptide-based drugs.

## Introduction

Natural products are secondary metabolites produced by living organisms—compounds not strictly required for growth and reproduction [1]. They are a productive source of drug leads, with approximately 40% of FDA-approved drugs derived from or inspired by them [2]. Among the most pharmacologically significant natural products are nonribosomal peptides (NRPs): this class includes the glycopeptide antibiotic vancomycin, the immunosuppressant cyclosporin A, the lipopeptide antibiotic daptomycin, and antifungal echinocandins, alongside numerous siderophores and cytostatic agents in clinical or preclinical use. NRPs are biosynthesized by nonribosomal peptide synthetases (NRPSs), large modular enzymes whose assembly-line biosynthetic logic enables remarkable structural and chemical diversity [1, 3, 4].

NRPSs adopt a strictly modular organization in which each module is responsible for the incorporation and functional modification of a single substrate—typically an amino acid, often a non-proteinogenic one. In the canonical case, a module consists of three core domains that are homologous across different NRPSs: an adenylation (A) domain, which selects and activates the cognate substrate as an aminoacyl-adenylate; a thiolation (T) domain (also called a peptidyl carrier protein, PCP), which tethers the activated substrate as a phosphopantetheine-linked thioester; and a condensation (C) domain, which catalyzes peptide bond formation between adjacent T-domain-tethered intermediates. Beyond these core domains, modules may carry optional modification domains that further diversify the final product, including epimerisation (E), cyclisation (Cy), methylation (MT), oxidation (Ox), reduction (Red), thioesterase (TE), and specialized starter and terminal condensation domains (C_start_, C_term_).

Among the three core domains, A-domains act as the central gatekeepers of substrate selection and thereby exert a decisive influence on the chemical identity of the final peptide product; engineering their specificity to incorporate non-native substrates has accordingly been a long-standing goal. Structural studies of the archetypal GrsA A-domain (PheA; PDB: 1AMU) revealed that ten conserved sequence motifs (A1–A10) cluster spatially around an active site embedded within a stretch of approximately 100 amino acids, and that substrate recognition is largely mediated by a subset of residues lining the binding pocket. Crystallographic and sequence analysis identified ten residues within this active site as the primary determinants of substrate specificity; this set is collectively termed the ‘Stachelhaus code’ [5, 6]. The tenth residue (position 517), a distal and near-invariant lysine, is omitted from predictive use, so the Stachelhaus code is represented here by its nine variable positions. A subsequent structural study extended this set to 34 residues by including all side chains within 8 Å of the bound substrate in the GrsA crystal structure [7]; this ‘8 Å code’ has since become the de facto standard feature set for computational A-domain substrate prediction.

The majority of computational tools for A-domain substrate prediction are built upon either the Stachelhaus code or the 8 Å code. NRPSpredictor2 [8], SANDPUMA [9], AdenPredictor [10], and PARAS [11] all represent A-domains by encoding the 34-residue set as physicochemical descriptor vectors, including AAindex-derived properties and *z*-scale features [12, 13]. In each case the residue set is defined by a distance criterion applied to a reference structure, and the selected positions serve as input features for downstream classifiers. Alternative codes have been proposed alongside the 8 Å shell, most targeting fungal A-domains specifically: a 15-residue code from a 6 Å shell in acyl:CoA synthetase homology models [14]; a 13-residue code from comparative modeling of siderophore synthetases [15]; two 17-residue codes based on evolutionary analysis [16] and crystal structure [17] respectively; and an 18-residue code derived from a 5 Å shell in AlphaFold [18] models of fungal A-domains [19] (Figure 1A). More recently, AlphaFold has allowed PARAS and AdenPredictor to supplement these sequence-derived descriptors with explicit geometric features of the predicted binding pocket.

**Fig 1.**
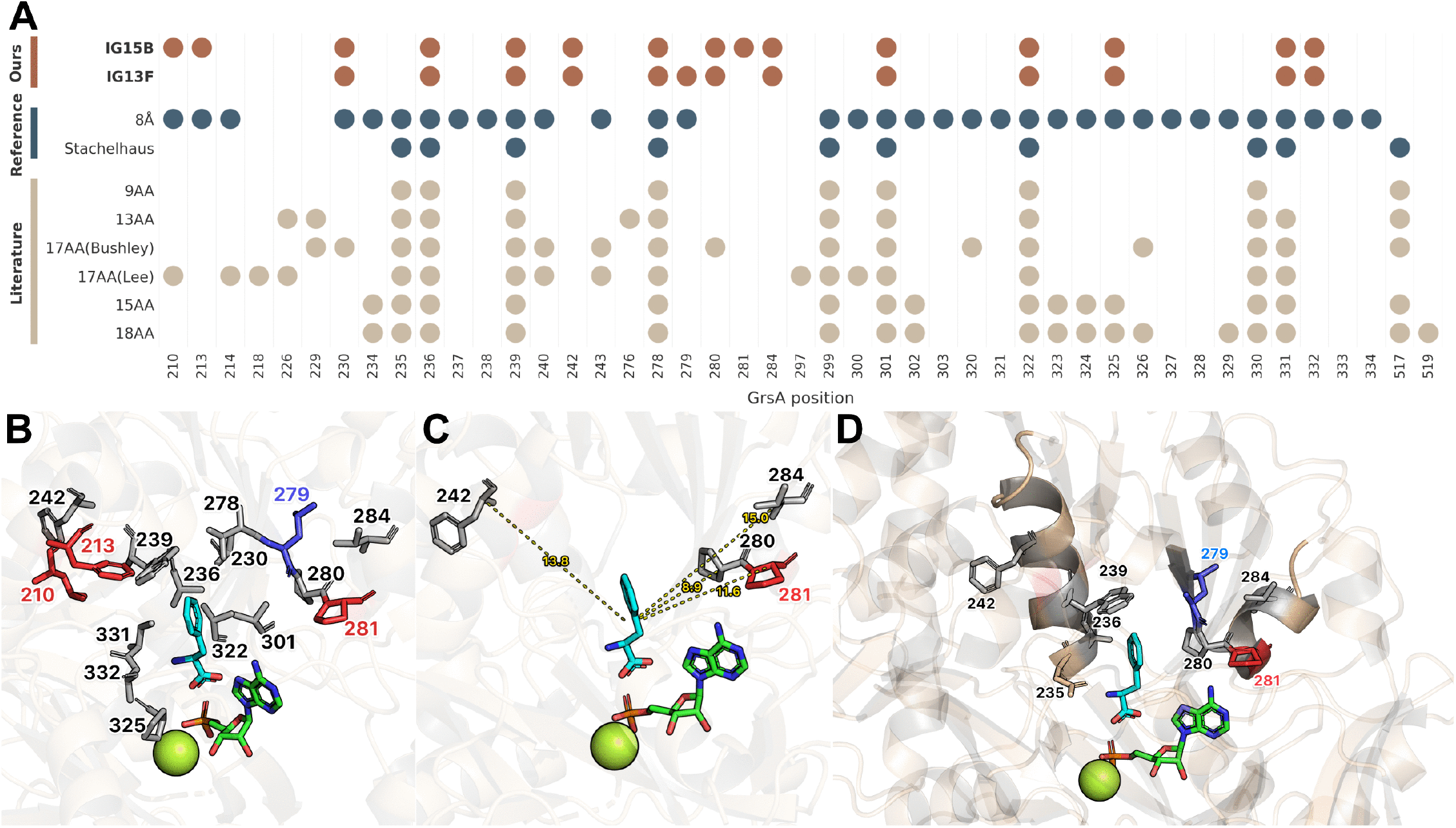
Substrate-specificity positions in NRPS A-domains. (**A**) Substrate-binding residue positions across A-domain codes. Circles mark residues included in each code. Rows are grouped into the codes proposed in this work (IG15B, IG13F), benchmark codes (8 Å pocket and Stachelhaus [5]), and literature codes for reference: 9AA [6], 13AA [15], 17AA(Bushley) [16], 17AA(Lee) [17], 15AA [14], and 18AA [19]. Numbering follows GrasA A (PheA; PDB: 1AMU). (**B**) Structural view of IG15B and IG13F residue sets within the A domain binding pocket, displayed alongside the co-crystallized substrate and AMP. Residues common to both sets are shown in gray, those unique to IG15B in red, and those unique to IG13F in blue. Substrate carbon atoms are rendered in cyan, while AMP carbon atoms and the Mg^2+^ ion are rendered in green. Residue numbering follows the GrsA_A convention. (**C**) Distances of novel positions 242, 280, 281, and 284 to the co-crystallized substrate. (**D**) Novel position 242 and classical Stachelhaus positions 235, 236, and 239 co-localize on the same *α*-helix; Position 280 marks the transition from a *β*-strand to an *α*-helix that harbors novel positions 281 and 284. Substrate carbon atoms are shown in cyan; AMP and Mg^2+^ ion in green.

NRPSTransformer [20] represents a departure from this framework entirely: by fine-tuning the protein language model ESM-2 [21] on full-length A-domain sequences, it achieves state-of-the-art performance without any predefined residue code, demonstrating that specificity information can be extracted directly from raw sequence context.

Despite this diversity of approaches, a fundamental limitation of the code-based tradition has gone largely unaddressed: all conventional residue sets—from the Stachelhaus code to the 8 Å shell and its variants—were defined by structural or geometric proximity to the substrate, rather than by any systematic assessment of which positions are informative for substrate specificity. Not all positions within the 8 Å shell contribute equally: highly conserved residues such as position 235 (aspartic acid in ~92% of sequences) reflect structural or catalytic constraints rather than variation that tracks substrate specificity, introducing redundancy into the feature set. Conversely, positions lying just beyond the distance threshold may carry genuine specificity information but are excluded by construction. Regardless of the sophistication of the model, a classifier can only exploit the residues that are given as input. As increasingly comprehensive A-domain sequence datasets have become available, it is now possible to revisit which residues govern substrate specificity from a purely data-driven, information-theoretic perspective—free of any prior assumption about spatial proximity to the substrate.

The contributions of this work are as follows:

- We assemble and curate a non-redundant A-domain dataset of 5,366 unique sequences (4,693 bacterial and 673 fungal) with substrate labels standardized against PubChem and ChEBI, providing a consistent basis for residue-level analysis.
- We introduce an information-theoretic framework that ranks alignment positions by their statistical association with substrate identity, without restricting the candidate pool to any predefined structural shell, thereby separating genuinely informative positions from conserved but uninformative ones.
- We identify two compact, kingdom-specific specificity-conferring codes—IG15B for bacterial and IG13F for fungal A-domains—which match or exceed the predictive performance of the 34-residue 8 Å code while using fewer than half its positions.
- We discover four positions (242, 280, 281, and 284) that lie outside both the Stachelhaus and 8 Å codes yet carry non-redundant specificity information, and show that they co-localize with classical determinants on two helices flanking the binding pocket, suggesting new candidate sites for A-domain engineering.

## Results

We assembled a non-redundant dataset of 5,366 A-domain sequences (4,693 bacterial, 673 fungal) spanning 430 substrates from three public resources, and applied a four-stage analysis pipeline (Figure 2): multiple sequence alignment against GrsA, position-level scoring by three information-theoretic measures—information gain (IG), mutual information (MI), and the *χ*^2^ statistic—and minimal-set selection by incrementally adding top-ranked positions as features to a random forest classifier evaluated by 5-fold cross-validation. A held-out benchmark of 150 bacterial and 130 fungal A-domains, excluded from the training data of every tool compared here, was used for all reported accuracies. Full dataset curation and procedural details are provided in Materials and Methods.

**Fig 2.**
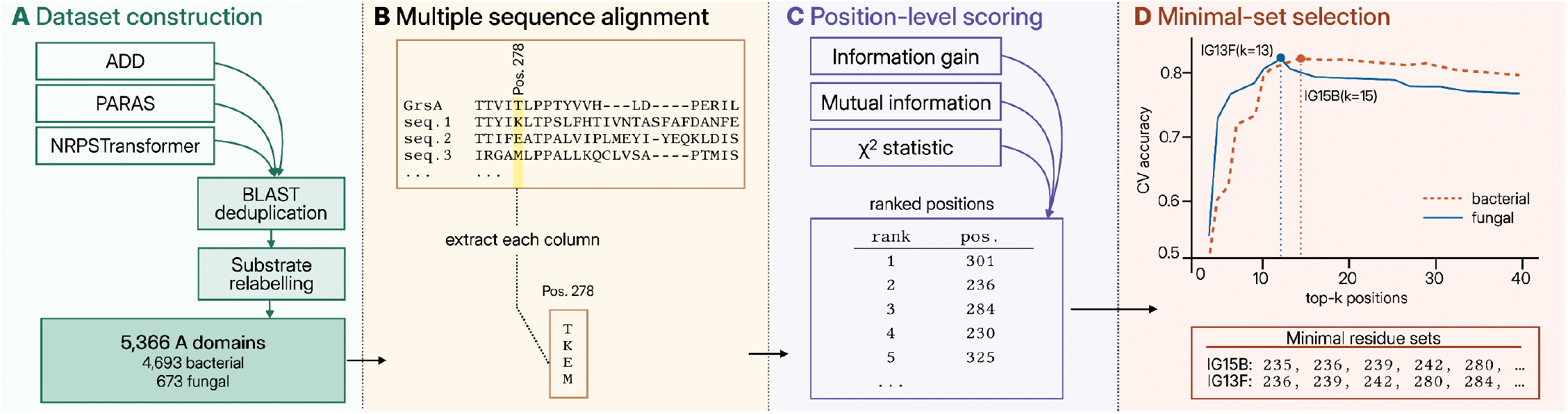
Overview of the data-driven framework for identifying substrate-specificity-conferring positions in NRPS A-domains. The pipeline proceeds through four stages from left to right. (**A**) *Dataset construction*: A-domain sequences are aggregated from three public resources (ADD, PARAS, and NRPSTransformer), deduplicated by BLAST at 100% sequence identity, and re-annotated against PubChem and ChEBI to standardize substrate labels, yielding 5,366 unique A-domains (4,693 bacterial and 673 fungal). (**B**) *Multiple sequence alignment* : All sequences are aligned with MUSCLE5 against GrasA_A as the reference, so that columns can be referenced by a common position index (Pos. 278 shown as an example). Each column is then extracted as a categorical feature describing the amino acid identity at that aligned position across the dataset. (**C**) *Position-level scoring* : Each alignment column is scored by three measures of statistical association with substrate identity—information gain (IG), mutual information (MI), and the *χ*^2^ statistic—and positions are ranked by their scores. (**D**) *Minimal-set selection*: Top-ranked positions are incrementally added as features for a random forest classifier, evaluated by 5-fold cross-validated accuracy on the bacterial (red, dashed) and fungal (blue, solid) training sets. The minimal residue set for each kingdom is defined as the smallest *k* at which accuracy reaches a local optimum, yielding IG15B (*k*=15) for bacterial A-domains and IG13F (*k*=13) for fungal A-domains.

### Information-theoretic scoring yields compact, kingdom-specific specificity codes

Among the three scoring measures, IG produced the highest cross-validated classification accuracy while requiring fewer positions than MI or *χ*^2^, and we therefore adopted it for the downstream analyses. For bacterial A-domains, the procedure converged on a set of 15 top-ranked positions (hereafter IG15B); for fungal A-domains, an analogous analysis identified 13 top-ranked positions (hereafter IG13F).

We compared IG15B and IG13F with the classical Stachelhaus code and the 8 Å code (Figure 1B). IG15B recapitulated six of the nine Stachelhaus positions and five further positions from the 8 Å code, while nominating four positions absent from both conventional codes (242, 280, 281, and 284). IG13F recovered the same six Stachelhaus positions and four further positions shared with the 8 Å code, while additionally selecting three positions absent from both conventional codes (242, 280, and 284).

### Novel positions cluster on two binding-pocket helices flanking the substrate

The four positions absent from both the Stachelhaus and 8 Å codes (242, 280, 281, and 284) were not distributed randomly across the binding pocket; instead, they co-localized with classical specificity-determining residues on two distinct *α*-helices (Figure 1C & D). The first helix harbours Stachelhaus positions 235, 236, and 239 alongside the novel position 242. The second secondary-structure element contains position 280, which marks the transition from a *β*-strand to an *α*-helix and is present in the 17AA(Bushley) code, together with the novel positions 281 and 284 on the adjacent *α*-helix. All three novel positions on the second helix (280, 281, 284) lay beyond the 8 Å distance threshold from the substrate (8.9 Å, 11.6 Å, and 15.0 Å, respectively), and position 242 on the first helix was similarly distal at 13.8 Å. These values nevertheless remain well below the mean centroid-to-centroid distance from the substrate averaged over all residues of GrasA_A (22.4 Å), indicating relative proximity of the newly identified positions, which can be regarded as next-to-the-second-shell residues.

### IG-derived codes match or exceed conventional codes with fewer positions

To assess how effectively each residue set captures substrate specificity, we conducted ablation experiments comparing classification accuracy across all codes, as well as their set-theoretic unions and intersections, using a random forest classifier trained on the full dataset and evaluated on the held-out benchmark comprising 150 bacterial and 130 fungal A-domains (Table 1).

**Table 1.** Classification accuracy of A-domain specificity-conferring codes evaluated on held-out benchmarks using a random forest classifier. |*S*| = code size; Δ = accuracy difference vs. the proposed kingdom-specific code (IG15B bacterial, IG13F fungal); Significance from Holm–Bonferroni-corrected McNemar test against the proposed code: ^∗^*p <* 0.05, ^∗∗^*p <* 0.01, ^∗∗∗^*p <* 0.001; n.s. = not significant. Proposed codes are shaded.

| (A) Bacterial benchmark ($N = 150$) | | | | | (B) Fungal benchmark ($N = 130$) | | | | |
| --- | --- | --- | --- | --- | --- | --- | --- | --- | --- |
| Residue set | $\vert S\vert $ | Acc. (%) | $\Delta$ | Sig. | Residue set | $\vert S\vert $ | Acc. (%) | $\Delta$ | Sig. |
| **IG15B (ours)** | 15 | 86.7 | — | — | **IG13F (ours)** | 13 | 69.2 | — | — |
| IG15B ∪ Stachelhaus | 18 | 88.7 | +2.0 | n.s. | 8 Å | 34 | 68.5 | −0.7 | n.s. |
| IG15B ∩ 8 Å | 11 | 86.7 | $\pm 0.0$ | n.s. | IG13F ∪ 8 Å | 37 | 66.9 | −2.3 | n.s. |
| 8 Å | 34 | 86.7 | $\pm 0.0$ | n.s. | IG13F ∪ Stachelhaus | 16 | 66.2 | −3.0 | n.s. |
| IG15B ∪ 8 Å | 38 | 86.0 | −0.7 | n.s. | Stachelhaus | 9 | 66.2 | −3.1 | n.s. |
| Stachelhaus | 9 | 85.3 | −1.3 | n.s. | IG13F ∩ 8 Å | 3 | 28.5 | −40.8 | *** |
| IG15B ∩ Stachelhaus | 6 | 82.0 | −4.7 | n.s. | IG13F ∩ Stachelhaus | 1 | 6.9 | −62.3 | *** |
| PARAS | — | 77.0 | −9.7 | ** |  |  |  |  |  |
| NRPSTransformer | — | 74.7 | −12.0 | *** |  |  |  |  |  |

For both bacterial and fungal A-domains, the IG-derived codes matched or exceeded the conventional 8 Å and Stachelhaus codes while using fewer than half as many positions, with the margin most pronounced for fungi. Their substantial overlap with the Stachelhaus code—six of its nine positions in the bacterial case—indicates that information-gain ranking recovers the most informative classical residues without any prior structural knowledge.

The set-theoretic comparisons reveal how this predictive signal is distributed. For bacterial A-domains, IG15B and the conventional codes share a common core: combining IG15B with either code changes accuracy only marginally, and their intersections retain nearly all of it, so the few non-overlapping residues carry only weak complementary signal. For fungal A-domains the picture inverts. IG13F overlaps minimally with the conventional codes, and restricting to either intersection collapses accuracy, while the corresponding unions fail to improve on IG13F alone. The predictive power of IG13F therefore resides in novel positions absent from both classical codes, where the additional conventional residues dilute rather than enhance the compact informative signal. PARAS could not be fairly benchmarked on the fungal set: its publicly released model is trained on essentially all of our fungal benchmark sequences and accordingly reproduces them at 100% accuracy by memorization, so we omit it from the fungal comparison (Table 1B). For reference, the original publication reports a fungal accuracy of 70.0% on its own held-out partition [11], which is not directly comparable to the matched evaluation reported here.

These results validate the information-theoretically derived codes on two fronts. First, they are at least as predictive as—and in the fungal case superior to—the established codes, while requiring substantially fewer positions, demonstrating that our approach distils the most informative residues from the alignment. Second, the substantial overlap between IG15B and the Stachelhaus code corroborates the biological relevance of our data-driven selection: the classical specificity-conferring residues are largely recovered without any prior structural knowledge, while the additionally identified positions provide further predictive gains. In the fungal context, where overlap with classical codes is minimal, the novel positions selected by IG13F are themselves the primary drivers of accuracy, underscoring the value of a kingdom-specific, information-theoretic approach to residue selection.

### Novel positions carry independent specificity information

To determine whether the novel positions carry genuinely independent specificity information or merely reflect co-evolution with adjacent residues from classical codes, we performed mutual information (MI) analysis on each *α*-helix separately, comparing each novel position against the classical Stachelhaus positions co-localized on the same helix (Figure 3). Specifically, the first helix harbors Stachelhaus positions 236 and 239 alongside the novel position 242, while the second region encompasses position 280, which lies at the junction between a *β*-strand and an *α*-helix and partially overlaps with the 17AA(Bushley) code, together with the novel positions 281 and 284 located on the adjacent *α*-helix. The logic of this test rests on two complementary quantities. The mutual information between two positions measures how redundant they are: if a novel position simply co-evolved with a classical neighbor, the two would be highly correlated and the novel position would carry no information the classical one lacks. The joint mutual information of a set of positions with the substrate class measures their combined predictive value: if the novel position were redundant, adding it to its classical neighbors would leave the joint value essentially unchanged, whereas an increase signals genuinely new information. A novel position is therefore non-redundant when it shows low inter-position MI with its classical neighbors yet raises the joint MI with substrate beyond what those neighbors provide alone.

**Fig 3.**
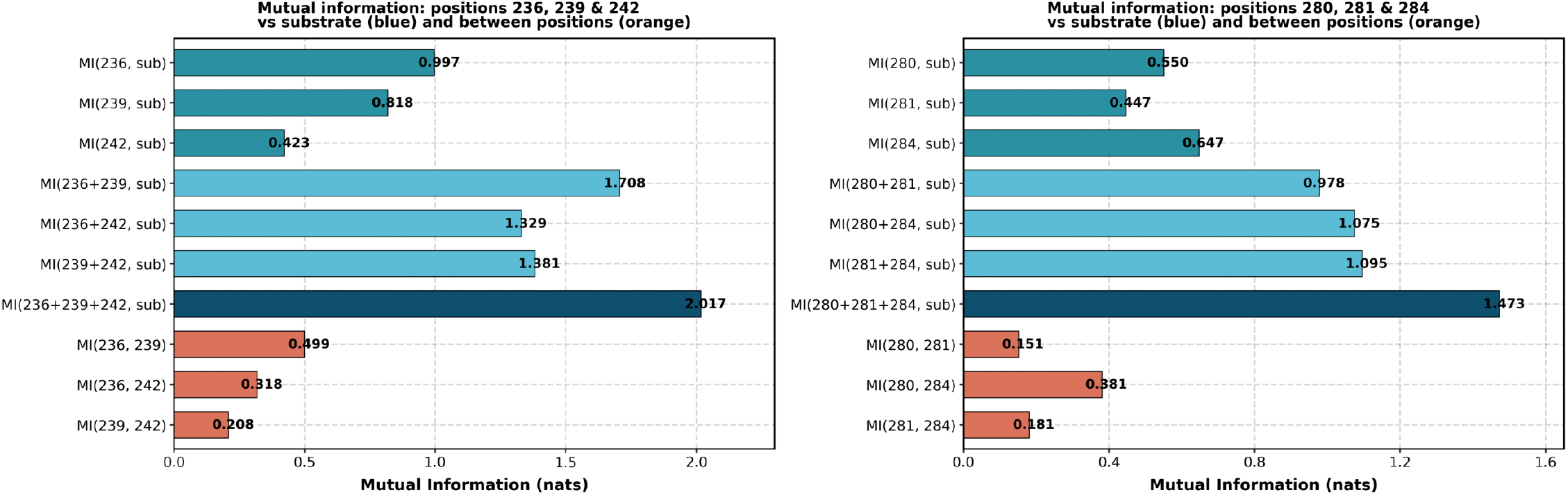
Mutual information (nats) among novel and classical specificity-conferring positions. Blue: marginal MI between each position and the substrate class; orange: inter-position MI between position pairs. Joint MI for each position set is reported in the text. Left: positions 236, 239, and 242; right: positions 280, 281, and 284.

In the first *α*-helix, the Stachelhaus positions 236 and 239 and the novel position 242 each have substantial marginal MI with the substrate class, but only moderate inter-position MI with one another (Figure 3), so they are not merely co-evolving duplicates. Decisively, their joint MI(236+239+242, substrate) = 2.017 nats exceeds the joint MI of any pair drawn from them—in particular that of the classical pair 236+239 alone—showing that position 242 supplies substrate information the two classical positions cannot.

The same pattern holds in the second region: positions 280, 281, and 284 each have substantial marginal MI with the substrate class but only low inter-position MI with one another (Figure 3), confirming they are largely independent. Their joint MI(280+281+284, substrate) = 1.473 nats again exceeds that of any pair, so the novel positions 281 and 284 contribute information beyond the partially classical position 280.

These results demonstrate that the novel positions identified by our information-theoretic approach are not statistical artifacts of co-evolution with residues from the classical codes. These mutual-information results also provide the statistical basis for the ablation experiment reported earlier (Table 1): the accuracy reduction observed when only the IG15B ∩ Stachelhaus intersection is retained reflects the loss of genuinely non-redundant specificity information carried by positions unique to each set—such as position 242—rather than the removal of redundant signal already captured by the shared core. Their exclusion from conventional codes is therefore a consequence of the distance-based selection criterion rather than an absence of functional relevance.

## Discussion

In this work we revisited which residues govern substrate specificity in NRPS A-domains from a purely data-driven perspective. Using information-theoretic measures to rank alignment positions by their statistical association with substrate identity, we identified two compact kingdom-specific codes—IG15B for bacterial and IG13F for fungal A-domains—that match or exceed the predictive accuracy of the classical Stachelhaus and 8 Å codes while using fewer than half the positions of the latter. Crucially, the same analysis identified four positions (242, 280, 281, and 284) that lie outside both conventional codes yet carry non-redundant specificity information. Our information-theoretic sets are designed to capture statistical associations rather than causal determinants, but their substantial overlap with historically established codes and independent corroboration of the novel positions (see below) supports their biological relevance.

The 34-residue 8 Å code was defined by including all residues within an 8 Å shell around the substrate in the GrsA crystal structure [7], and by design encompasses both first-shell residues directly contacting the substrate and second-shell residues shaping the binding pocket. This inclusive approach introduces redundancy: many positions are structurally or catalytically essential but display minimal variation across substrates, contributing little information for classification. Position 235 illustrates this limitation: included in both 8Å and Stachelhaus codes, it encodes aspartic acid in 92% of sequences, reflecting its conserved role in stabilizing the substrate *α*-amino group. We note that this residue is not entirely uninformative: its variation can distinguish A-domains activating *β*-amino acids from those activating canonical *α*-amino acids, and in this sense it carries genuine specificity information. Within the predominantly *α*-amino-acid substrate space considered here, however, its near invariance limits its informative power. Information-theoretic ranking naturally down-weights such highly conserved residues, yielding more compact and less redundant residue sets.

The four novel positions identified here are not statistical artifacts of co-evolution with classical residues. On each binding-pocket helix, the joint mutual information between the novel position and its co-localized Stachelhaus neighbors exceeds any pairwise combination (Figure 3), confirming that the novel positions contribute independent specificity information. Position 280 is further corroborated by independent structural evidence from fungal ferrichrome synthetases: it lies within *β*-Strand 2 (T275–P280), one of four sequence fragments that collectively define the walls of the substrate-binding pocket in the GrsA A-domain crystal structure [16], and it marks the transition between a *β*-strand and an *α*-helix that together form one wall of the pocket. Its side chain is positioned to influence the spatial dimensions of the cavity, contributing to substrate specificity through steric rather than purely electrostatic means. Their absence from conventional codes is therefore a consequence of the distance-based selection criteria underlying these codes, rather than an absence of functional relevance—a limitation that information-theoretic analysis is well-suited to address.

A complementary perspective on the value of these residue sets is provided by comparison with state-of-the-art classifiers. Existing tools—PARAS/PARASECT [11] and NRPSTransformer [20]—rely on considerably more elaborate machinery: PARAS and PARASECT incorporate AlphaFold-predicted structures [18] to extract geometric and spatial features of the binding pocket, while NRPSTransformer fine-tunes the protein language model ESM-2 [21] on raw A-domain sequences. On the same held-out benchmark (Table 1), our random forest classifiers built on IG15B and IG13F achieved comparable or higher accuracy with substantially fewer features. This is not strictly a head-to-head comparison—our classifier was trained and evaluated on a substrate space restricted to those present in the benchmark, whereas existing tools contend with the full open-set prediction problem—but the result suggests that feature selection may matter at least as much as model capacity for current A-domain substrate prediction.

Our approach offers several advantages that we believe account for its competitive performance. First, by ranking alignment positions according to their information content with respect to substrate identity, our method discriminates between residues that are merely conserved (and thus uninformative for classification, as in position 235) and those whose variation truly tracks substrate class. This naturally prunes redundancy from the feature space. Second, by not constraining the candidate pool to the canonical 8 Å shell, our approach is able to identify informative positions, such as 242, 280, 281, and 284, that lie beyond direct contact distance with the substrate but nonetheless contribute non-redundant predictive signal (Figure 3). Existing tools, however sophisticated their downstream models, draw their input from within the conventional 34-residue set and may therefore have limited access to such positions; the information available to a classifier is ultimately shaped by the residues it considers. Third, the resulting residue sets are compact and interpretable: unlike embeddings derived from protein language models or geometric features extracted from predicted structures, each position selected by IG15B and IG13F can be inspected, mapped onto the binding pocket, and reasoned about biologically. These properties suggest that feature selection may be at least as important as model capacity in current A-domain substrate prediction, and that the residues identified here could complement existing methods rather than replace them.

Beyond substrate prediction, the residue sets identified in this work may also offer guidance for A-domain engineering. Rational redesign of A-domain specificity has long been a central goal in NRPS engineering, yet it is frequently constrained by an incomplete understanding of which residues to mutate. Substitutions guided solely by the Stachelhaus code or the 8 Å shell often yield limited or unpredictable changes in substrate preference, in part because the residues considered may not exhaust those that contribute to selectivity. By highlighting positions such as 242, 280, 281, and 284—which carry independent specificity information yet have remained outside conventional codes—our analysis offers an additional set of candidate sites for mutagenesis and combinatorial library design. We anticipate that incorporating these positions into engineering campaigns, alongside the classical specificity-conferring residues, could expand the accessible sequence space for redesigning A-domain specificity and ultimately facilitate the production of novel nonribosomal peptides.

These findings suggest that the conventional specificity-conferring residue sets, while biologically meaningful, are neither minimal nor exhaustive. The positions identified here provide a more compact and information-rich representation of A-domain specificity and may complement, rather than replace, existing prediction tools. More broadly, the same information-theoretic strategy is applicable to any ligand–protein dataset in which aligned protein sequences are paired with discrete ligand labels, and we hope this work will encourage a broader reassessment of which residues are considered in future studies of A-domain specificity.

## Materials and methods

### Dataset assembly and curation

A-domain sequences were compiled from three public resources: the ADD database^1^, the PARAS dataset [11], and the NRPSTransformer dataset [20]. Because these resources are derived from overlapping repositories such as MIBiG [22], redundant entries were expected, and the merged collection was deduplicated using BLAST [23]: two sequences were considered identical when their pairwise alignment showed 100% sequence identity, so that even single-amino-acid variants—which may confer distinct substrate specificities—were preserved as unique entries. Identical sequences originating from different NRPS proteins were treated as a single data point representing the same functional A-domain. After deduplication, the dataset comprised 5,366 unique A-domain sequences (4,693 bacterial, 673 fungal).

Substrate labels were re-annotated and standardized against PubChem [24] and ChEBI [25]. Chirality at the side-chain level was merged during relabeling (e.g., 3S-hydroxyasparagine and 3R-hydroxyasparagine were both annotated as 3-hydroxyasparagine), while stereochemistry at the *α*-carbon was retained (l-alanine and d-alanine remained distinct labels). The harmonized dataset spans 430 unique substrates.

Two filtering steps were applied before training: sequences with multiple substrate annotations were excluded to ensure unambiguous sequence–substrate assignments, and the training substrate vocabulary was restricted to substrates represented in the benchmark to allow direct comparison with existing tools on a consistent substrate space. The benchmark sets comprised 150 bacterial A-domains covering 49 substrates and 130 fungal A-domains covering 23 substrates, curated from the NRPSTransformer benchmark [20] and further refined for this work; no benchmark sequence appears in the training data of any tool compared in this study. After filtering, the training set comprised 4,420 bacterial A-domains (45 substrates) and 524 fungal A-domains (24 substrates).

### Multiple sequence alignment

A multiple sequence alignment (MSA) of all A-domain sequences in the dataset was constructed using MUSCLE5 [26] with the `-super5` mode, which is optimized for large-scale alignments with high accuracy. The GrsA(UniProt Entry: P0C061) was included as a reference sequence to facilitate positional comparison across all alignments.

### Position scoring

Three statistical methods—information gain (IG), mutual information (MI), and *χ*^2^ statistic—were employed to identify the most discriminative positions across all aligned A-domain sequence positions with respect to substrate specificity.

#### Notation

Let term *t*_*i,a*_ denote the co-occurrence of amino acid *a* at position *i*, where *i* ∈ {1, 2, …, *L*} (*L* is the length of the multiple sequence alignment) and *a* ∈ A (the set of 20 standard amino acids). The substrate specificity, denoted *s*, represents the category (e.g., a specific amino acid or analog). The dataset comprises *N* sequences, with probabilities estimated empirically from the training set. Let 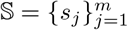 be the set of *m* substrate categories.

#### Information gain (IG)

IG quantifies the reduction in substrate specificity uncertainty given knowledge of term *t*_*i,a*_:

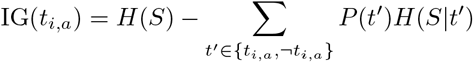

where the entropy of substrates is:

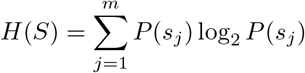

and the conditional entropy is:

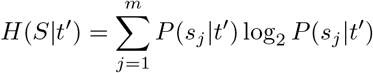

Here, *P* (*s*_*j*_) is the prior probability of substrate *s*_*j*_, *P* (*t*^′^) is the probability of term presence or absence, and *P* (*s*_*j*_|*t*^′^) is the conditional probability.

#### Mutual information (MI)

MI measures the information shared between term *t*_*i,a*_ and substrate *s*:

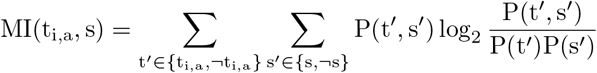

where *P* (*t*^′^, *s*^′^) is the joint probability, and *P* (*t*^′^), *P* (*s*^′^) are marginal probabilities. MI is zero for independent *t*_*i,a*_ and *s*. Global relevance is assessed via:

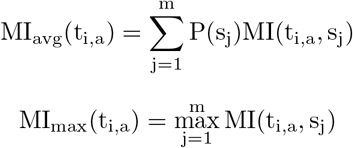

#### *χ*^2^ statistic

The *χ*^2^ statistic tests dependence between *t*_*i,a*_ and *s* using a 2 × 2 contingency table, where *N*_11_ is the number of sequences with both *t*_*i,a*_ and *s, N*_10_ with *t*_*i,a*_ but not *s, N*_01_ with *s* but not *t*_*i,a*_, *N*_00_ with neither, and *N* the total sequences:

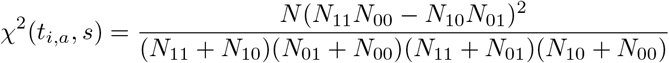

A zero *χ*^2^ indicates independence. Global scores are:

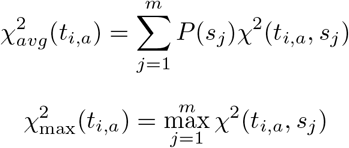

#### Position-Specific Scoring

For each position *i* in the aligned sequences, a position-specific score is computed by maximizing over all amino acids *a* ∈ A:

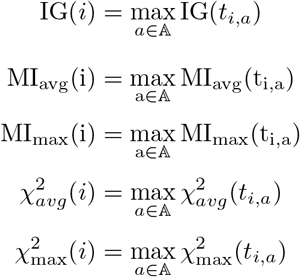

The maximum score is selected because substrate specificity in A-domains is often driven by a specific amino acid at a critical position, making the highest-scoring amino acid the most discriminative for that position. This approach ensures that positions with the greatest influence on substrate recognition are prioritized.

### Minimal residue set selection and classifier training

A minimal set of alignment positions sufficient to specify A-domain substrate specificity was identified for bacterial and fungal A-domains separately, since the two kingdoms display distinct sequence patterns. For each kingdom, alignment positions were ranked in descending order by their position-specific scores (defined above) and used to construct cumulative feature sets; amino acid identities at each selected position were one-hot encoded as model inputs.

For each candidate size *k*, a random forest classifier with 500 trees was trained on the top-*k* ranked positions, with five independent models trained under different random seeds. Five-fold cross-validation was used to identify the value of *k* at which mean accuracy reached a local optimum, and the resulting model was then evaluated on the held-out benchmark. Accuracy was defined at the substrate-set level: a prediction was considered correct if the predicted label matched any substrate in the annotated substrate set for that A-domain.

For bacterial A-domains, classifiers were trained on bacterial sequences only. For fungal A-domains, training on fungal sequences alone yielded broad, poorly defined accuracy curves without a clear peak at small *k*. To obtain a more stable and interpretable minimal residue set, classifiers for fungal A-domains were retrained on a combined dataset of bacterial and fungal sequences, with performance evaluated only on the fungal benchmark. This mixed-training strategy substantially sharpened the accuracy peak and yielded more compact residue sets.

## Footnotes

1 https://github.com/Chaotic-algorithm/ADDdatabase

## Notes

### Competing Interest Statement

The authors have declared no competing interest.

## References

1. Demain AL, Fang A. In: Fiechter A, editor. The Natural Functions of Secondary Metabolites. Berlin, Heidelberg: Springer Berlin Heidelberg; 2000. p. 1–39. Available from: https://doi.org/10.1007/3-540-44964-7_1. doi:10.1007/3-540-44964-7_1.

2. Newman DJ, Cragg GM. Natural Products as Sources of New Drugs over the Nearly Four Decades from 01/1981 to 09/ 2019;83(3):770–803. Available from: http://doi.org/10.1021/acs.jnatprod.9b01285. doi:10.1021/acs.jnatprod.9b01285.

3. Walsh CT. Polyketide and Nonribosomal Peptide Antibiotics: Modularity and Versatility. Science. 2004;303(5665):1805–10. Available from: https://www.science.org/doi/abs/10.1126/science.1094318. arXiv: https://www.science.org/doi/pdf/10.1126/science.1094318. doi:10.1126/science.1094318.

4. Sieber SA, Marahiel MA. Molecular Mechanisms Underlying Nonribosomal Peptide Synthesis:â€‰ Approaches to New Antibiotics. Chemical Reviews. 2005;105(2):715–38. PMID: 15700962. Available from: https://doi.org/10.1021/cr0301191. arXiv:https://doi.org/10.1021/cr0301191. doi:10.1021/cr0301191.

5. Stachelhaus T, Mootz HD, Marahiel MA. The Specificity-Conferring Code of Adenylation Domains in Nonribosomal Peptide Synthetases. Chemistry & Biology. 1999 Aug;6(8):493–505. doi:10.1016/S1074-5521(99)80082-9.

6. Challis GL, Ravel J, Townsend CA. Predictive, Structure-Based Model of Amino Acid Recognition by Nonribosomal Peptide Synthetase Adenylation Domains. Chemistry & Biology. 2000 Mar;7(3):211–24. doi:10.1016/s1074-5521(00)00091-0.

7. Rausch C, Weber T, Kohlbacher O, Wohlleben W, Huson DH. Specificity Prediction of Adenylation Domains in Nonribosomal Peptide Synthetases (NRPS) Using Transductive Support Vector Machines (TSVMs);33(18):5799–808. Available from: https://doi.org/10.1093/nar/gki885. doi:10.1093/nar/gki885.

8. Röttig M, Medema MH, Blin K, Weber T, Rausch C, Kohlbacher O. NRPSpredictor2—a Web Server for Predicting NRPS Adenylation Domain Specificity. Nucleic Acids Research. 2011 Jul;39(suppl 2):W362–7. doi:10.1093/nar/gkr323.

9. Chevrette MG, Aicheler F, Kohlbacher O, Currie CR, Medema MH. SANDPUMA: Ensemble Predictions of Nonribosomal Peptide Chemistry Reveal Biosynthetic Diversity across Actinobacteria. Bioinformatics. 2017 Oct;33(20):3202–10. doi:10.1093/bioinformatics/btx400.

10. Mongia M, Baral R, Adduri A, Yan D, Liu Y, Bian Y, et al. AdenPredictor: Accurate Prediction of the Adenylation Domain Specificity of Nonribosomal Peptide Biosynthetic Gene Clusters in Microbial Genomes. Bioinformatics. 2023 Jun;39(Supplement 1):i40–6. doi:10.1093/bioinformatics/btad235.

11. Terlouw BR, Huang C, Meijer D, Cediel-Becerra JDD, He R, Rothe ML, et al. PARAS: High-Accuracy Machine-Learning of Substrate Specificities in Nonribosomal Peptide Synthetases. Bioinformatics; 2025. doi:10.1101/2025.01.08.631717.

12. Wold S, Eriksson L, Hellberg S, Jonsson J, Sjöström M, Skagerberg B, et al. Principal Property Values for Six Non-Natural Amino Acids and Their Application to a Structure–Activity Relationship for Oxytocin Peptide Analogues. Canadian Journal of Chemistry. 1987 Aug;65(8):1814–20. doi:10.1139/v87-305.

13. Kawashima S, Pokarowski P, Pokarowska M, Kolinski A, Katayama T, Kanehisa M. AAindex: Amino Acid Index Database, Progress Report 2008. Nucleic Acids Research. 2008 Jan;36(suppl 1):D202–5. doi:10.1093/nar/gkm998.

14. Khurana P, Gokhale RS, Mohanty D. Genome Scale Prediction of Substrate Specificity for Acyl Adenylate Superfamily of Enzymes Based on Active Site Residue Profiles;11(1):57. Available from: https://bmcbioinformatics.biomedcentral.com/articles/10.1186/1471-2105-11-57. doi:10.1186/1471-2105-11-57.

15. Schwecke T, Göttling K, Durek P, Dueñas I, Käufer NF, Zock-Emmenthal S, et al. Nonribosomal Peptide Synthesis in Schizosaccharomyces Pombe and the Architectures of Ferrichrome-Type Siderophore Synthetases in Fungi;7(4):612–22. arXiv:16502473. doi:10.1002/cbic.200500301.

16. Bushley KE, Ripoll DR, Turgeon BG. Module Evolution and Substrate Specificity of Fungal Nonribosomal Peptide Synthetases Involved in Siderophore Biosynthesis;8:328. arXiv:19055762. doi:10.1186/1471-2148-8-328.

17. Lee TV, Johnson LJ, Johnson RD, Koulman A, Lane GA, Lott JS, et al. Structure of a Eukaryotic Nonribosomal Peptide Synthetase Adenylation Domain That Activates a Large Hydroxamate Amino Acid in Siderophore Biosynthesis *;285(4):2415–27. Available from: https://www.jbc.org/article/S0021-9258(19)63736-1/abstract. arXiv:19923209. doi:10.1074/jbc.M109.071324.

18. Jumper J, Evans R, Pritzel A, Green T, Figurnov M, Ronneberger O, et al. Highly Accurate Protein Structure Prediction with AlphaFold;596(7873):583–9. Available from: https://www.nature.com/articles/s41586-021-03819-2. doi:10.1038/s41586-021-03819-2.

19. Heard S, Winter J. Structure-Guided Investigation of Fungal Adenylation Domain Substrate Selectivity. ChemRxiv. 2023;2023(1206). Available from: https://chemrxiv.org/doi/abs/10.26434/chemrxiv-2023-c24qz. arXiv: https://chemrxiv.org/doi/pdf/10.26434/chemrxiv-2023-c24qz. doi:10.26434/chemrxiv-2023-c24qz.

20. Zhang Z, Zhou Y, Xie S, Liu RZ, Huang Z, Saravana Kumar P, et al. NRPStransformer, an Accurate Adenylation Domain Specificity Prediction Algorithm for Genome Mining of Nonribosomal Peptides. Journal of the American Chemical Society. 2025 Aug:jacs.5c08076. doi:10.1021/jacs.5c08076.

21. Lin Z, Akin H, Rao R, Hie B, Zhu Z, Lu W, et al. Evolutionary-Scale Prediction of Atomic-Level Protein Structure with a Language Model. Science. 2023 Mar;379(6637):1123–30. doi:10.1126/science.ade2574.

22. Terlouw BR, Blin K, Navarro-Muñoz JC, Avalon NE, Chevrette MG, Egbert S, et al. MIBiG 3.0: A Community-Driven Effort to Annotate Experimentally Validated Biosynthetic Gene Clusters. Nucleic Acids Research. 2023 Jan;51(D1):D603–10. doi:10.1093/nar/gkac1049.

23. Altschul SF, Gish W, Miller W, Myers EW, Lipman DJ. Basic Local Alignment Search Tool;215(3):403–10. arXiv:2231712. doi:10.1016/S0022-2836(05)80360-2.

24. Kim S, Chen J, Cheng T, Gindulyte A, He J, He S, et al. PubChem 2025 update. Nucleic Acids Research. 2024 11;53(D1):D1516–25. Available from: https://doi.org/10.1093/nar/gkae1059. arXiv: https://academic.oup.com/nar/article-pdf/53/D1/D1516/60743708/gkae1059.pdf. doi:10.1093/nar/gkae1059.

25. Malik A, Arsalan M, Moreno C, Mosquera J, Félix E, Kizilören T, et al. ChEBI: re-engineered for a sustainable future. Nucleic Acids Research. 2025 11;54(D1):D1768–78. Available from: https://doi.org/10.1093/nar/gkaf1271. arXiv: https://academic.oup.com/nar/article-pdf/54/D1/D1768/65596350/gkaf1271.pdf. doi:10.1093/nar/gkaf1271.

26. Edgar RC. Muscle5: High-accuracy Alignment Ensembles Enable Unbiased Assessments of Sequence Homology and Phylogeny. Nature Communications. 2022 Nov;13(1):6968. doi:10.1038/s41467-022-34630-w.

